# Transient accumulation and bidirectional movement of KIF13B in primary cilia

**DOI:** 10.1101/2021.07.09.451768

**Authors:** Alice Dupont Juhl, Zeinab Anvarian, Stefanie Kuhns, Julia Berges, Jens S. Andersen, Daniel Wüstner, Lotte B. Pedersen

**Affiliations:** Department of Biochemistry and Molecular Biology, University of Southern Denmark, Campusvej 55, DK-5230 Odense M, Denmark; Department of Biology, University of Copenhagen, Universitetsparken 13, DK-2100 Copenhagen Ø, Denmark; Department of Biomedicine, Facultad Ciencias Experimentales, Universidad Francisco de Vitoria, Ctra. Pozuelo-Majadahonda Km. 1.800, 28223 Pozuelo de Alarcón (Madrid), Spain

## Abstract

Primary cilia are microtubule-based sensory organelles whose assembly and function rely on the conserved bidirectional intraflagellar transport (IFT) system, which is powered by anterograde kinesin-2 and retrograde cytoplasmic dynein 2 motors. Nematodes additionally employ a cell type-specific kinesin-3 motor, KLP-6, which moves within cilia independently of IFT and regulates ciliary content and function. Here we provide evidence that a KLP-6 homolog, KIF13B, undergoes bursts of bidirectional movement within primary cilia of cultured immortalized human retinal pigment epithelial (hTERT-RPE1) cells. Anterograde and retrograde intraciliary velocities of KIF13B were similar to those of IFT (IFT172-eGFP), but intraciliary movement of KIF13B required its own motor domain and appeared to be cell-type specific. Our work provides the first demonstration of motor-driven, intraciliary movement by a vertebrate kinesin other than kinesin-2 motors.

## Introduction

Primary cilia are microtubule-based sensory organelles found on the surface of almost all animal cells and are pivotal for regulating diverse signaling pathways such as Sonic hedgehog (Shh) and polycystin-1/polycystin-2-mediated signaling (Anvarian et al., 2019). Cilia are compartmentalized organelles whose assembly, maintenance and function depend on the conserved intraflagellar transport (IFT) machinery that via kinesin-2 and cytoplasmic dynein 2 motors, respectively, brings IFT-A and IFT-B complexes with associated ciliary cargoes into and out of the organelle (Pedersen and Rosenbaum, 2008, Taschner and Lorentzen, 2016). Ciliary composition and function are additionally regulated by the ciliary transition zone (Garcia-Gonzalo and Reiter, 2016), vesicular trafficking and endocytosis of ciliary components (Pedersen et al., 2016, Blacque et al., 2017), and release of extracellular vesicles (EVs) from cilia or the periciliary membrane compartment (Wood and Rosenbaum, 2015, Akella and Barr, 2021).

Two types of ciliary kinesin-2 motors have been described: heterotrimeric kinesin-2 (KIF3A/KIF3B/KAP3 in vertebrates) that functions as the canonical anterograde IFT motor, and homodimeric kinesin-2 (KIF17 in vertebrates) that functions as an accessory anterograde IFT motor in some cilia (Pedersen and Rosenbaum, 2008, Prevo et al., 2017). Several additional kinesins were shown to localize to and function at cilia, most of which regulate ciliary assembly and length by affecting microtubule dynamics (Reilly and Benmerah, 2019). A notable exception is the *Caenorhabditis elegans* kinesin-3 motor KLP-6, which moves independently of conventional IFT within male-specific sensory cilia, and negatively affects the velocity of the homodimeric kinesin-2 motor OSM-3/KIF17 (Morsci and Barr, 2011). KLP-6 also regulates ciliary targeting and EV-mediated release of polycystin-2 to control male mating behavior (Peden and Barr, 2005, Wang et al., 2014). Ciliary EVs act as signal-carrying entities in e.g. *C. elegans* and *Chlamydomonas* (Wang et al., 2014, Wood et al., 2013, Luxmi et al., 2019) and may rid the organelle of unwanted material in vertebrates (Nager et al., 2017, Phua et al., 2017). The precise mechanisms underlying release of EVs from cilia are unclear, but in addition to KLP-6, other factors implicated in this process include the BBSome, a retrograde IFT membrane cargo adaptor (Lechtreck, 2015, Nager et al., 2017, Akella et al., 2020), actin polymerization at the ciliary EV release site (Nager et al., 2017), and axonemal tubulin post translational modification (Akella and Barr, 2021). It is unknown if vertebrate kinesin-3 motors move within primary cilia and promote ciliary EV release, and to date KLP-6 is the only kinesin apart from IFT kinesin-2 motors with demonstrated intraciliary motility.

We previously showed that a KLP-6 homolog, KIF13B, localizes to primary cilia in mouse fibroblasts and immortalized human retinal pigment epithelial (hTERT-RPE1) cells, and that its depletion in these cells causes ciliary accumulation of the cholesterol-binding membrane protein CAV1 and impaired Shh signaling (Schou et al., 2017). The molecular basis for this phenotype was not clear, and it was unknown if KIF13B actually moves within cilia. Here, we show by live cell imaging combined with image analysis and simulations that KIF13B undergoes bursts of IFT-like bidirectional movement within primary cilia of hTERT-RPE1 cells. Anterograde and retrograde intraciliary velocities of KIF13B were similar to those of IFT (IFT172-eGFP), but intraciliary movement of KIF13B required its own motor domain and appeared to be cell-type specific. Occasionally, EV-like release of KIF13B from the cilium tip was observed. Our results provide the first demonstration of intraciliary movement by a vertebrate kinesin other than IFT kinesin-2 motors, and of transient/burst-like kinesin-driven motility within cilia of any organism.

## Results and discussion

To investigate if KIF13B moves inside cilia we expressed a previously validated eGFP-KIF13B fusion protein (Schou et al., 2017, Serra-Marques et al., 2020) in hTERT-RPE1 cells stably expressing a red fluorescent ciliary membrane marker, SMO-tRFP (Lu et al., 2015), and serum starved cells to induce ciliogenesis prior to analysis by confocal live cell imaging. In all cells examined (n=54), eGFP-KIF13B was strongly concentrated at the ciliary base, and 15 of these additionally displayed rapid and transient accumulation of eGFP-KIF13B within the cilium itself (Figure 1; Movie 1; Movie 2; Table 1). Moreover, among the 15 cells showing intraciliary movement of eGFP-KIF13B, 2 cells additionally exhibited EV-like release of eGFP-KIF13B from the ciliary tip (Figure 1A; Movie 1; Table 1). This rare phenomenon could be a side-effect of the ciliary SMO-tRFP overexpression as we did not observe it in hTERT-RPE1 cells expressing endogenously mScarlet-tagged ARL13B to mark cilia (Figure S1D; Movie 3; Table 1), although we cannot exclude that EV-like tip release of eGFP-KIF13B might occur in the latter cells on rare occasions and/or under specific physiological conditions. Importantly, in 16% of the transfected ARL13B-mScarlet cells (n=38) eGFP-KIF13B did exhibit rapid and transient accumulation within cilia, as observed in the SMO-tRFP cells (Figure S1D; Movie 3; Table 1), supporting that this behavior of eGFP-KIF13B was not a consequence of the ciliary membrane marker used. The proportion of transfected live hTERT-RPE1 cells displaying eGFP-KIF13B within cilia (16-37%; Table 1) is in line with our previous observations in fixed hTERT-RPE1 cells, where we detected eGFP-KIF13B in cilia of 23% of the transfected cells (Schou et al., 2017). Moreover, in the latter study, cells were pre-extracted with detergent containing AMP-PNP prior to fixation and immunofluorescence microscopy, supporting that eGFP-KIF13B is bound to axonemal microtubules within cilia (Schou et al., 2017).

**Figure 1.**
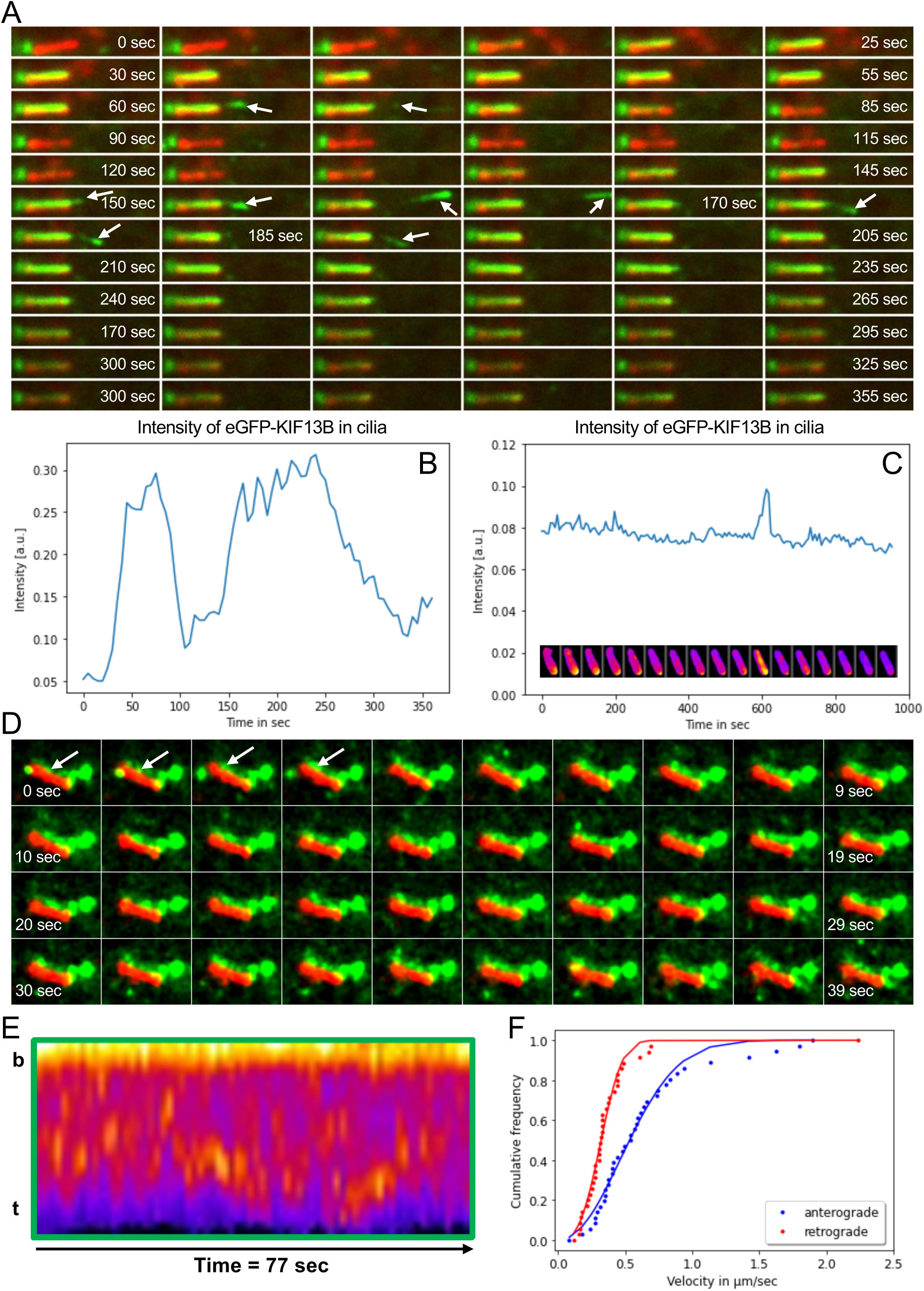
Transient bidirectional movement and EV-like tip release of eGFP-KIF13B in primary cilia of hTERT-RPE1 cells. (**A**) eGFP-KIF13B (green) is transiently moving into cilia, marked with SMO-tRFP (red). In less than one minute, eGFP-KIF13B accumulates in cilia, and some is released from the cilia tip (arrows) between 70-82 sec and 155-185 sec of the video sequence. Based on Movie 1. (**B**) Measurement of eGFP-KIF13B fluorescence intensity in cilia over time showing entry into cilia in repeated bursts. (**C**) Example of similar analysis as in (B) with only one burst of eGFP-KIF13B observed (under the graph are example snapshots aligned with the curve). (**D**) Time-lapse sequence of another cell acquired with higher time resolution, i.e., an interval time of 1.1 sec, and showing intraciliary movement of eGFP-KIF13B. Note that red and green images are slightly off-set in (A) and (D), to better visualize movement of eGFP-KIF13B relative to SMO-tRFP. Based on Movie 2. (**E**) Kymograph of such a high-time resolution sequence showing eGFP-KIF13B intraciliary movement (see also Figure S1E that includes lines). **b** and **t**, cilia base and tip. (**F**) Weibull distribution function of the form *v*, was fitted by a Weibull distribution function of the form *y*(*ν*) =1 - exp [−(*α* · *ν*)^*β*^] with α=1.58 and β=2.11 for anterograde and α=2.76 and β=2.94 for retrograde transport, respectively, suggests a rather broad distribution of eGFP-KIF13B velocities.

**Table 1.** Overview of live cell imaging results in hTERT-RPE1 (RPE1) cells, focusing on localization of GFP-tagged KIFs to the cilium-centrosome axis. n, number of live cells imaged per condition.

| Cell type/Condition | Fusion protein | n | Number of cells with KIF fusion protein localizing to: |  | Comment |
| --- | --- | --- | --- | --- | --- |
|  |  |  | Basal body | Cilium+basal body |  |
| RPE1-SMO-tRFP | eGFP-KIF13B | 54 | 39<br>(72%) | 15<br>(28%) | Burst-like movement in cilia. EV-like release from cilia observed in 2 cases. |
| RPE1-SMO-tRFP + Ciliobrevin | eGFP-KIF13B | 59 | 38<br>(64%) | 16<br>(27%) | Burst-like movement in 1 cilium. |
| RPE1-SMO-tRFP + DMSO | eGFP-KIF13B | 19 | 12<br>(63%) | 7<br>(37%) | Burst-like movement in 4 cilia. |
| RPE1-SMO-tRFP | eGFP-KIF13BΔM | 27 | 19<br>(70%) | 7<br>(26%) | No burst-like movement in cilia. |
| RPE1-SMO-tRFP | eGFP-KIF13A | 18 | 0 | 1<br>(6%) | - |
| RPE1-SMO-tRFP | GFP-KIF17 | 2 | 0 | 2<br>(100%) | Stable localization at the cilium tip. |
| RPE1-ARL13B-mScarlet | GFP-KIF17 | 14 | 0 | 14<br>(100%) | Stable localization at the cilium tip. |
| RPE1-ARL13B-mScarlet | eGFP-KIF13B | 38 | 32<br>(84%) | 6<br>(16%) | Burst-like movement in cilia. |
| IMCD3- mCherry-Arl13b | eGFP-KIF13B | 10 | 10<br>(100%) | 0<br>(0%) | - |
| IMCD3- mCherry-Arl13b + Purmorphamine | eGFP-KIF13B | 16 | 16<br>(100%) | 0<br>(0%) | - |

Kymograph analysis indicated that eGFP-KIF13B moves bidirectionally within cilia at 0.64 ± 0.07 μm/s in the anterograde direction (mean ± sem of n=37 measurements from 5 cells) and 0.39 ± 0.06 μm/s in the retrograde direction (mean ± sem of n=36 measurements from 5 cells) (Figure 1E and F, Figure S1C). This is within the range of velocities reported for anterograde and retrograde IFT in mammalian cells (Williams et al., 2014) and for anterograde velocity of KLP-6 in *C. elegans* CEM cilia (Morsci and Barr, 2011), but slower than the reported anterograde speed for KIF13B moving on cytoplasmic microtubules in e.g. HeLa cells (2-3 μm/s; (Serra-Marques et al., 2020)). Since kinesin velocity is influenced by multiple factors, including tubulin posttranslational modifications (Sirajuddin et al., 2014) and interaction with other motors (Snow et al., 2004, Morsci and Barr, 2011, Prevo et al., 2015, Milic et al., 2017), the unique structure and composition of the cilium may account for the slower KIF13B velocity in this compartment compared to the cell body. Nevertheless, kymographs indicated that eGFP-KIF13B moves progressively in both directions within cilia without slowing or acceleration and without prolonged pauses at the ciliary tip. From the cumulative histogram of velocities, we find that anterograde movement is slightly faster than retrograde movement (Figure 1E and F). The cumulative histogram of velocities, *v*, was fitted by a Weibull distribution function of the form *y*(*ν*) = 1 - exp [−(*α* · *ν*)^*β*^] with α=1.58 and β=2.11 for anterograde and α=2.76 and β=2.94 for retrograde transport, respectively, suggesting a rather broad distribution of velocities (Figure 1F). Image-based simulations of IFT indicate that movement of eGFP-KIF13B inside cilia with such velocities over the entire cilia length of < 5 µm would result in oscillation frequencies for bidirectional movement of about 30 sec (Figure S2). The observed periodicity of intensity oscillations of eGFP-KIF13B in cilia was much slower; typically, between 100-500 sec (Fig. 1B and Movie 1, 2). This can be inferred from the example shown in Fig. 1A and B and is supported by an analysis of power spectra of this and similar intensity profiles (Fig. S1A, B). Such dynamic behavior likely reflects the import and export kinetics of KIF13B in primary cilia combined with frequent switching of directions. It is comparable to the avalanche-like injection of IFT particles into cilia, observed previously (Ludington et al., 2013), indicating that eGFP-KIF13B moves frequently up and down in cilia before eventually being exported from the cilia base into the cytosol or, in rare cases, released from the cilium tip.

In *C. elegans* amphid/phasmid sensory cilia homodimeric (OSM-3) and heterotrimeric kinesin-2 motors function in coordination to mediate anterograde IFT in the axoneme middle segment, whereas OSM-3 alone mediates anterograde IFT in the distal segment (Snow et al., 2004). In male-specific *C. elegans* cilia the kinesin-3 motor KLP-6 moves independently of OSM-3 and heterotrimeric kinesin-2 IFT motors, but KLP-6 appears to slow down the OSM-3 motor (Morsci and Barr, 2011). To investigate if eGFP-KIF13B intraciliary motility is coordinated with conventional IFT, we first compared its dynamics to that of the IFT-B2 complex component IFT172 (Taschner et al., 2016), which we endogenously tagged with eGFP at its C-terminus (IFT172-eGFP) in the hTERT-RPE1 cell line. We found that IFT172-eGFP moves bidirectionally along cilia in distinct spots, whose paths are clearly discernable in the video sequences and kymographs (Figure 2A-D and Movie 4). Anterograde and retrograde movement of IFT172-eGFP was occasionally observed in parallel in the same cilium (Figure 2C, D), and IFT172-eGFP was constantly present in cilia in contrast to the periodic ciliary import/export observed for eGFP-KIF13B. Kymograph analysis indicated that IFT172-eGFP moves with an average velocity of 0.53 ± 0.02 μm/s in the anterograde direction (mean ± sem of n=75 measurements from 5 cells) and of 0.32 ± 0.02 μm/s in the retrograde direction (mean ± sem of n=50 measurements from 5 cells). The cumulative histogram of velocities of IFT172-eGFP was also fitted to a Weibull distribution function with α=1.58 and β=2.11 for anterograde transport, and α=2.83 and β=2.74 for retrograde transport, respectively (Figure 2E). Both the mean values and the distribution function of measured velocities of IFT172-eGFP coincide very closely with the measured intraciliary velocities of eGFP-KIF13B in the hTERT-RPE1 SMO-tRFP cells. These results suggest that intraciliary movement of KIF13B is closely coordinated with conventional IFT. To further sustain this conclusion, we attempted to visualize intraciliary eGFP-KIF13B movement simultaneously with mCherry-IFT88, expressed stably in hTERT-RPE1 cells, but this was not feasible as cells expressing both fusion proteins failed to produce cilia. Therefore, we next carried out live cell imaging analysis in the hTERT-RPE1 SMO-tRFP cells treated with the dynein inhibitor Ciliobrevin D (Firestone et al., 2012), which was previously used to inhibit IFT in mammalian primary cilia (Ye et al., 2013). These analyses showed that eGFP-KIF13B localized within primary cilia of 27% of the transfected, Ciliobrevin D-treated cells (n=59), but no oscillatory movement of eGFP-KIF13B within cilia was detected under these conditions (Figure 3A, B; Table 1; Movie 5). Instead, eGFP-KIF13B very slowly entered cilia, as shown in Fig. 3A and Movie 5, or it was exported in a time course of several minutes (not shown). Since Ciliobrevin D treatment leads to accumulation of stalled IFT trains on axonemal microtubules (Ye et al., 2013), we cannot conclude from these experiments whether eGFP-KIF13B is being moved by IFT as the “roadblock” effect of stalled IFT trains could impair eGFP-KIF13B intraciliary movement indirectly. To further investigate possible coordination of eGFP-KIF13B intraciliary movement with conventional IFT, quantitative live cell imaging analysis was additionally performed using a truncated version of KIF13B lacking the motor domain (Lamason et al., 2010), hereafter referred to as motorless eGFP-KIF13B. This analysis demonstrated that motorless eGFP-KIF13B enters cilia in 26% of the transfected cells (n=27), but its fluorescence intensity was much lower in cilia than at the basal body, and bursts of intraciliary movement and ciliary tip release were not observed (Figure 3C, D; Table 1; Movie 6). In contrast, similar analysis of a motorless version of homodimeric kinesin-2 KIF17 in cilia of mammalian olfactory sensory neurons showed that it moved along the entire axoneme by associating with heterotrimeric kinesin-2 motors (Williams et al., 2014). We conclude that intraciliary motility of KIF13B depends on its own motor domain and is closely coordinated with conventional IFT.

**Figure 2.**
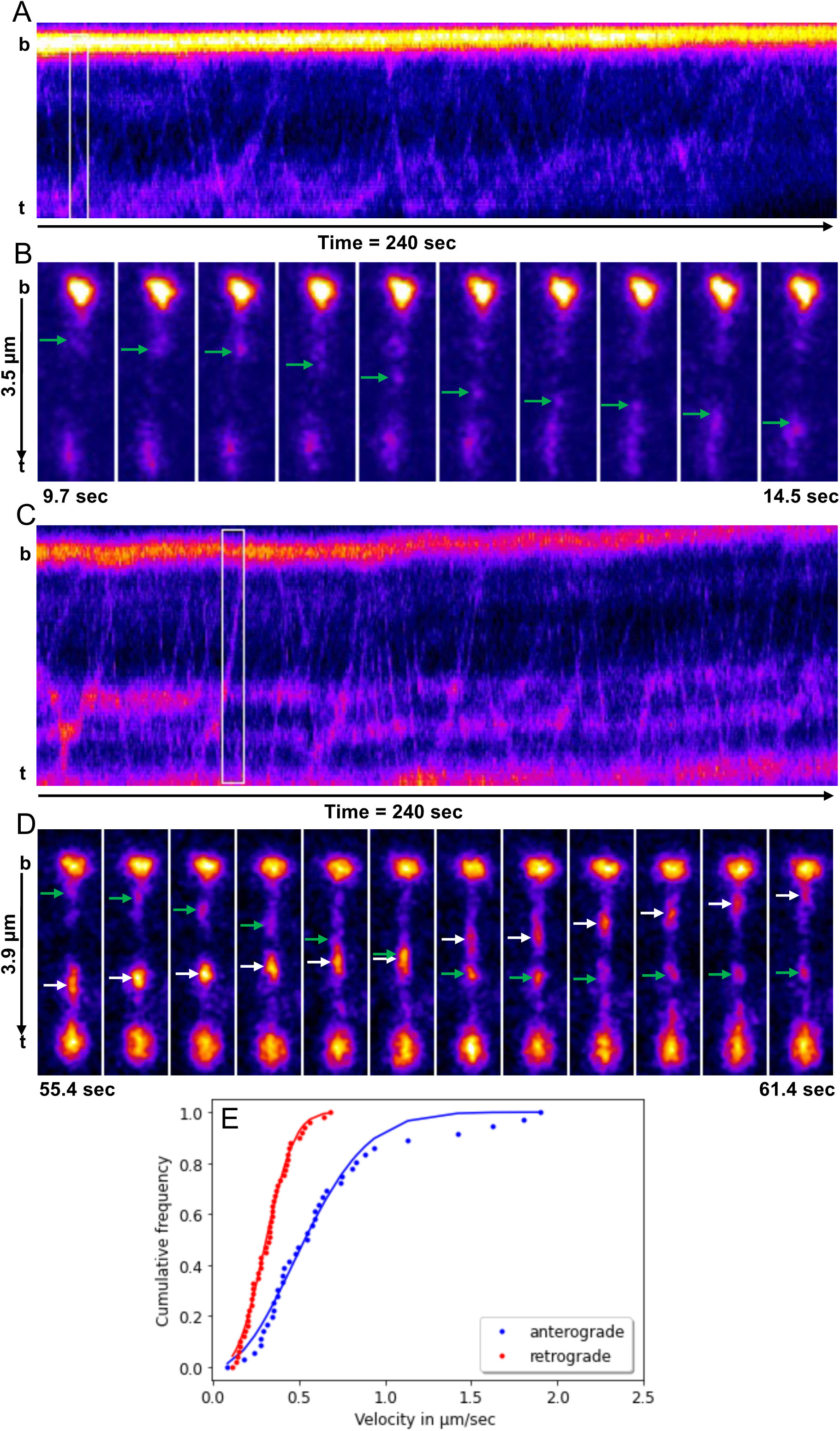
Live cell imaging analysis of IFT172-eGFP in cilia. (**A, C**) Kymographs from two hTERT-RPE1 cells stably expressing IFT172-eGFP and imaged with high temporal resolution (1.9 frames/second) at a confocal microscope with temperature control. Based on Movie 4. (**B**) Zoom of the kymograph shown in panel A (white box) with anterograde movement from the cilia base (**b**) to tip (**t**, green arrows). (**D**) Zoom of the kymograph shown in panel C (white box) with anterograde movement from the cilia base (**b**) to tip (**t**, green arrows) crossing with retrogradely moving IFT172-eGFP (white arrows in D). (**E**) Weibull distribution function of the form *v*, was fitted by a Weibull distribution function of the form *y*(*ν*) =1 - exp [−(*α* · *ν*)^*β*^] with α=1.58 and β=2.11 for anterograde and α=2.83 and β=2.74 for retrograde transport, respectively.

**Figure 3.**
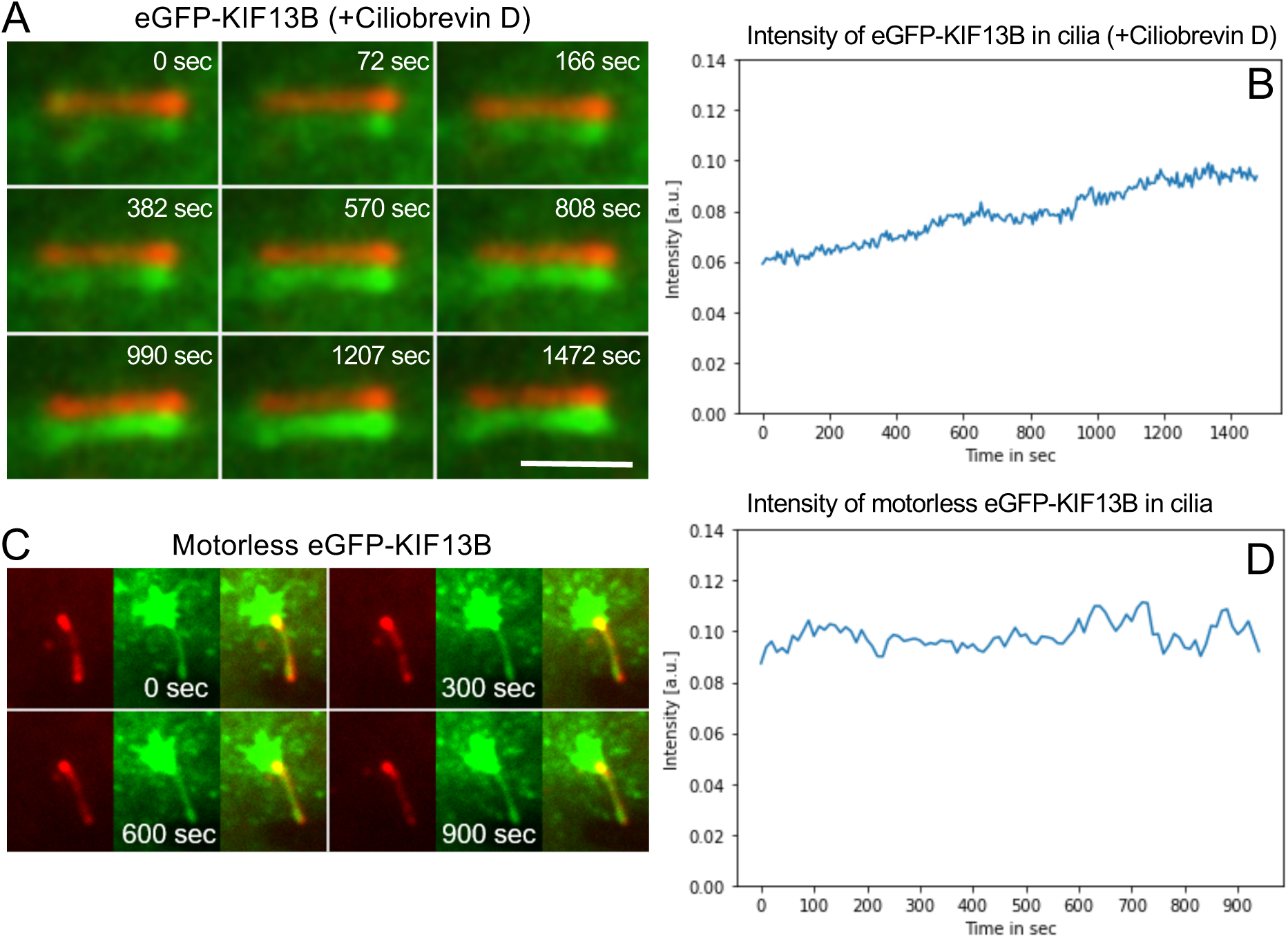
Ciliary movement of eGFP-KIF13B requires its own motor domain. (**A**) Snapshots of Movie 5 showing eGFP-KIF13B cilia localization in SMO-tRFP cells treated with 10 μM Ciliobrevin D for 1-6 hours. (**B**) Measurement of eGFP-KIF13B fluorescence intensity in cilia of Ciliobrevin D treated cells as a function of time, based on Movie 5 and similar. (**C**) Snapshots of Movie 6 showing motorless eGFP-KIF13B in SMO-tRFP (red) cells accumulating around the ciliary base with some localizing stably to cilia over time (n=25 cells). (**D**) Measurement of motorless eGFP-KIF13B fluorescence intensity in cilia over time, based on Movie 6 and similar.

Next, we asked whether other kinesin-3 motors or known ciliary motors exhibit the same type of burst-like intraciliary motility as we observed for eGFP-KIF13B. The human genome codes for seven different kinesin-3 motors (Hirokawa et al., 2010) of which the KIF13 subgroup, comprised by KIF13B and its proximal paralog KIF13A, displays the highest sequence homology to the ciliary *C. elegans* KLP-6 kinesin-3 motor (Schou et al., 2017). When we analyzed eGFP-KIF13A movement in live hTERT-RPE1 SMO-tRFP cells, we only detected eGFP-KIF13A in the cilium of a single transfected cell out of 18 cells analyzed (6%; Figure S3A; Table 1), which is in agreement with our previous results using fixed hTERT-RPE1 cells (Schou et al., 2017). In contrast, GFP-tagged KIF17, which functions as an accessory IFT kinesin-2 motor in some vertebrate cell types (Jenkins et al., 2006, Insinna et al., 2008, Williams et al., 2014), was stably localized at the ciliary tip in all cells examined (Figure S3B; Table 1), as expected (Dishinger et al., 2010). Thus, amongst the motors tested, burst-like intraciliary motility was only observed for KIF13B.

To investigate if intraciliary movement of KIF13B is cell-type specific, as is the case for *C. elegans* KLP-6 (Peden and Barr, 2005), we next performed live cell imaging analysis of eGFP-KIF13B in ciliated mouse inner medullary collecting duct 3 (IMCD3) cells stably expressing mCherry-Arl13b. We found that eGFP-KIF13B localizes to the base of, but not within, cilia of these cells (Figure S3C; Table 1). It is unclear why eGFP-KIF13B fails to localize to cilia in mCherry-ARL13B expressing IMCD3 cells; notably in preliminary experiments we observed that mNeonGreen-tagged KIF13B localizes to and moves within cilia of mouse cortical collecting duct (mCCD) cells (data not shown), suggesting that intraciliary movement of KIF13B is not limited to hTERT-RPE1 cells.

In summary, our results demonstrate that eGFP-KIF13B transiently enters cilia and moves bidirectionally within the organelle at speeds reminiscent of IFT. Occasionally we observed EV-like release of eGFP-KIF13B from the ciliary tip. This phenomenon could be a side-effect of the ciliary membrane marker used for imaging, but we cannot exclude that physiologically relevant release of KIF13B from cilia might occur under some circumstances, e.g. in response to physical (Wang et al., 2020) or chemical (Nager et al., 2017, Phua et al., 2017) stimuli. Intraciliary movement of eGFP-KIF13B requires its own motor domain and seems to be cell-type specific, as reported for KLP-6 in *C. elegans* (Morsci and Barr, 2011). Since KIF13B is a plus end-directed motor and moves bidirectionally within cilia at similar velocities as IFT172, its retrograde movement must be mediated via association of KIF13B with the dynein-2 driven retrograde IFT machinery. Anterograde movement of KIF13B within cilia is likely powered by its own intrinsic motor activity yet coordinated with kinesin-2 motors/conventional IFT. We also cannot exclude that the KIF13B motor domain is required for interaction with the IFT machinery. In future studies it will be interesting to further dissect the coordination of KIF13B intraciliary movement with IFT, e.g. using single molecule imaging technologies, and to determine the ciliary cargoes of KIF13B.

## Materials and methods

### Plasmids, mammalian cell culture, stable cell lines and transfection

The following previously described plasmids were used: eGFP-KIF13B (Asaba et al., 2003), motorless eGFP-KIF13B (Lamason et al., 2010), eGFP-KIF13A (Schou et al., 2017), GFP-KIF17 (Jaulin and Kreitzer, 2010), pcDNA3.1 (+) (Thermo Fisher Scientific), pCAG-enAsCas12a (Addgene plasmid # 107941) (Kleinstiver et al., 2019), pMaCTag-Z12 (Addgene plasmid # 120055) and pMaCTag-P05 (Addgene plasmid # 120016) (Fueller et al., 2020). Plasmids were propagated in *Eschericia coli* DH10B cells using standard procedures and purified using NucleoBond Xtra Midi EF Kit (Macherey-Nagel).

hTERT-RPE1 cells stably expressing SMO-tRFP were generously provided by Christopher J. Westlake and have been described previously (Lu et al., 2015). To establish the hTERT-RPE1 cell lines with ARL13B-mScarlet or IFT172-eGFP tagged at the endogenous locus, we used a CRISPR/Cas12a-assisted PCR tagging approach (Fueller et al., 2020). Briefly, repair donor templates were generated by PCR with gene specific primers and the plasmids pMaCTag-Z12 or pMaCTag-P05. The PCR template cassette together with the enAsCas12a expression plasmid were electroporated into hTERT-RPE1 cells using the Neon Transfection System (Thermo Fisher), and cells were selected with zeocin or puromycin for 2 weeks. Isolated single cell clones were characterized by live cell imaging, PCR and Sanger sequencing. The hTERT-RPE1 SMO-tRFP cells were cultured in Dulbecco’s Modified Eagle Medium (DMEM; Thermo Fisher Scientific) supplemented with 10% fetal bovine serum (FBS; Thermo Fisher Scientific), 1% penicillin-streptomycin (Thermo Fisher Scientific), and 10 µg/mL Hygromycin B. hTERT-RPE1 ARL13B-mScarlet and hTERT-RPE1 IFT72-eGFP cells were cultured in DMEM/F12 medium (Thermo Fischer Scientific) supplemented with 10% FBS (Gibco), 1 % Glutamax (Gibco) and 1% penicillin-streptomycin. hTERT-RPE1 cells stably expressing mCherry-IFT88 were generated from lentiviral transduction using plasmids described in (Campeau et al., 2009). The IFT88 ORF was first amplified using PCR (forward primer: 5’-CCGGTACCATGATGCAAAATGTGCACCTGGC-3’; reverse primer: 5’-CCGCGGCCGCTTATTCTGGAAGCAAATCATCTCCTAAT-3’) from pCAG2-mChe-IFT88 (Kobayashi et al., 2021), generously provided by Kazuhisa Nakayama. PCR products were cloned into the Gateway system-compatible pENTR-mCherry-C1. Subsequently, pENTR-mCherry-IFT88 was recombined with pCDH-EF1a-Gateway-IRES-BLAST plasmid with LR recombination (Invitrogen/Thermo Fisher Scientific). Pools of hTERT-RPE1 cells stably expressing mCherry-IFT88 were selected with Blasticidin (Sigma). For Ciliobrevin D experiments, culture medium was exchanged with M1 medium (150 mM NaCl_2_, 5 mM KCL, 1 mM CaCl_2_, 1 mM MgCl_2_, 5 mM glucose, and 20 mM HEPES) containing 10 µM Ciliobrevin D (Merck) in DMSO. Imaging was started after 1h, and control cells treated with a similar volume of DMSO were analyzed in parallel.

Mouse IMCD3 cells stably expressing mCherry-ARL13B were a generous gift from Peter Gorilak and Vladimir Varga at Laboratory of Cell Motility, Institute of Molecular Genetics of the Czech Academy of Sciences, Prague, Czech Republic. These were cultured in DMEM/F12 medium (Thermo Fischer Scientific) containing 10% heat inactivated FBS and 1% penicillin-streptomycin. For the imaging experiments, cells were seeded on glass-bottom dishes 35/22 mm (HBSt-3522) from Willco Wells or on 35 mm microscope dishes with glass bottom from MatTek (P35G-1.5–50-C). Cells were transfected with KIF-encoding plasmid and empty vector pcDNA3.1 (+) (1 µg total DNA) using 2 µl Lipofectamine 3000 Reagent and 2 µl P3000 (Thermo Fisher Scientific). hTERT-RPE1 cells stably expressing SMO-tRFP were transfected a day after seeding and IMCD3 cells expressing mCherry-Arl13b were reverse transfected at the time of seeding. To induce ciliogenesis, cells were cultured overnight (16 hr) in a medium free of FBS and antibiotics. For hTERT-RPE1 cells, serum starvation started 2 hr post transfection and for IMCD3 cells ca. 16 hr post transfection. To activate the Shh pathway and promote SMO ciliary entrance in IMCD3 cells, the cells were serum starved with DMEM/F12 medium containing 2 µM purmorphamine (Sigma, cat# SML0868).

Cell lines were routinely authenticated and checked for contamination.

### Fluorescent live cell imaging

Live cell imaging was carried out on several confocal microscopes, all temperature (37°C) and CO_2_ (5%) regulated in a humidity chamber. These included a fully motorized Olympus IX83 inverted microscope equipped with a spinning disc (Yokogawa) and a Hamatsu ORCA-Flash 4.0 digital camera (C13440); a motorized inverted Nikon-Andor spinning disk microscope equipped with CSU-X1 spinning disk (Yokogawa), and/or laser launcher, Okolab microscope stage incubator, and a perfect focus system (PFS) (at DaMBIC, University of Southern Denmark). Both 60x and 100x Numerical Aperture (NA) 1.4 oil objectives (Olympus) were used. The 488 and 561 laser lines were used for imaging eGFP and tRFP, respectively. The time-lapse sequences were obtained with time intervals ranging from 5 to 10 sec. For laser scanning confocal microscopy at DaMBIC, University of Southern Denmark (SDU), a Nikon Ti-2, A1 LFO confocal microscope with a Plan Apo λ 100 x NA 1.4 oil objective with the following excitation/emission settings was used: 488/525 nm for eGFP and 561/595 nm for tRFP and mScarlet. The interval for the time-lapse sequences was ranging from 0.5 to 10 sec. All live cell imaging was carried out in DMEM or M1 medium containing 150 mM NaCl_2_, 5 mM KCL, 1 mM CaCl_2_, 1 mM MgCl_2_, 5 mM glucose, and 20 mM HEPES. During imaging experiments, we carefully selected those cells that had a low expression level of eGFP-KIF13B (or other kinesin fusion proteins) but could still be visualized with our microscope. In most cases, cells overexpressing eGFP-KIF13B lose their primary cilia because of the negative effect of this protein on the cilia length, as reported in (Schou et al., 2017).

### Image analysis and simulations

All image analysis and simulations were carried out in ImageJ (https://imagej.nih.gov/ij/), either using existing plugins, or implemented as self-developed Macro scripts (available from the authors upon request). Subsequent time series analysis was implemented as Jupyter notebooks in Python (https://jupyter.org/). First, cilia intensity of eGFP-tagged KIF13B (wild-type or motorless mutant) was isolated based on the red fluorescent marker protein SMO-tRFP. For that, a rectangular region of interest (ROI) was defined around the red cilia marker, and an intensity threshold was applied to the SMO-tRFP fluorescence in cilia weighted by the mean intensity of the red marker in the entire ROI. This procedure accounted for eventual photobleaching and secured a stable cilia area segmentation. A binary mask comprising the cilia area with intensities of either 0 or 1 was generated and multiplied with the corresponding green image series of the eGFP-tagged KIF13B. Integrated fluorescence intensity of eGFP-KIF13B was measured from videos with long acquisition time (i.e. 5-6 sec, acquired at a spinning disk confocal system) and normalized to the total green fluorescence in the rectangular ROI. To determine periodicities in cilia location dynamics of eGFP-KIF13B, the extracted intensity time series was transformed into frequency space using the Fast Fourier transform function of the scipy library in Python. Time-lapse sequences with shorter acquisition time (i.e. 1.1.sec intervals) were treated identically, but in addition to calculating integrated intensities, a Kymograph analysis was carried out. For that, the segmented cilia were first spatially aligned using a rigid body registration procedure. A 3-pixel wide line (straight or segmented) was used and the kymograph calculated using the Multikymograph plugin to Image J developed by Drs. A.Seitz and J. Rietdorf. Straight lines were identified and velocities of eGFP-KIF13B were calculated using the accompanying Macro script based on the known pixel size and interval time. Cumulative histograms of velocities measured in the anterograde and retrograde direction were calculated and fitted to a Weibull function as described in the Results and discussion section. Image simulations of cilia movement were implemented as ImageJ Macro with a fixed cilia length of 5 µm, a simulated pixel size of either 0.05 or 0.1 µm and a frame rate of 1 frame per sec (1 Hz). A variable number of consecutive motors was translated towards the cilia tip with a constant speed. A stochastic velocity component was added from a uniform distribution, comprising about 0-10% of the total velocity to account for the stochastic nature of motor movement. Simulated images were convoluted with a Gaussian filter of width equal to 0.15 or 0.25 µm and Gaussian noise was added to account for the blurring by the microscope optics and the camera read out noise, respectively.

## Acknowledgements

We thank Drs Christopher J. Westlake, Valdimir Varga, Athar Chisthi, Joel Pomerantz, Geri Kreitzer, Kazuhisa Nakayama, Michael Knop, Keith Joung and Benjamin Kleinstiver for reagents, and Søren Johansen for technical assistance. Supported by grants from Novo Nordisk Foundation (NNF18OC0053024, NNF15OC0016886 and NNF14OC0011535), Danish Cancer Society (R146-A9590) and Hartmann Fonden (A31662) to LBP. JB was supported by an Eramus+ traineeship from the European Commision. DW acknowledges funding from the Villum Foundation (Grant No. 35865). JSA and SK acknowledge support from Independent Research Fund Denmark (Grant No. 8021-00425B). We thank Danish Molecular Biomedical Imaging Center, University of Southern Denmark, for use of imaging equipment, supported by Novo Nordisk Foundation (NNF18SA0032928).

