## Supplementary figures and movie legends for "Transient accumulation and bidirectional movement of KIF13B in primary cilia"

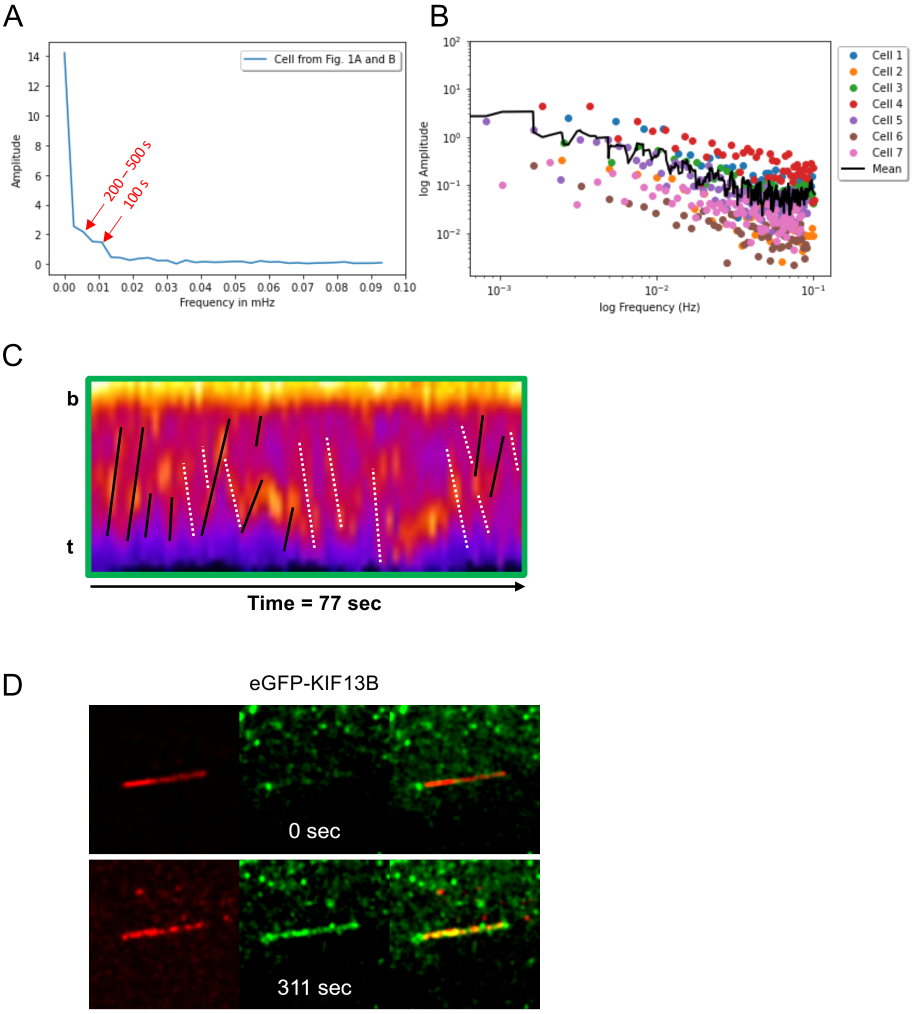


**Figure S1. Live cell imaging analysis of eGFP-KIF13B in hTERT-RPE1 SMO-tRFP and ARL13B-mScarlet cells. Related to Figure 1.** (**A**) Fourier transformation (power spectrum) of the data in Figure 1B with main periodicities indicated. (**B**) Log-log plot of power spectra from several (n=7) analyzed cells with mean shown as black line. (**C**) Kymograph of high-time resolution sequences, similar to Figure 1E, showing lines for anterograde (straight lines) and retrograde (dashed lines) eGFP-KIF13B intraciliary movement. (**D**) Example snapshots of movie showing burst-like intraciliary movement of eGFP-KIF13B in hTERT-RPE1 ARL13B-mScarlet cells. A total of 38 cells were imaged, of which 6 showed burst-like intraciliary movement of eGFP-KIF13B. The movie was acquired with a speed of 0.9

frames/second. See the corresponding Movie 3.

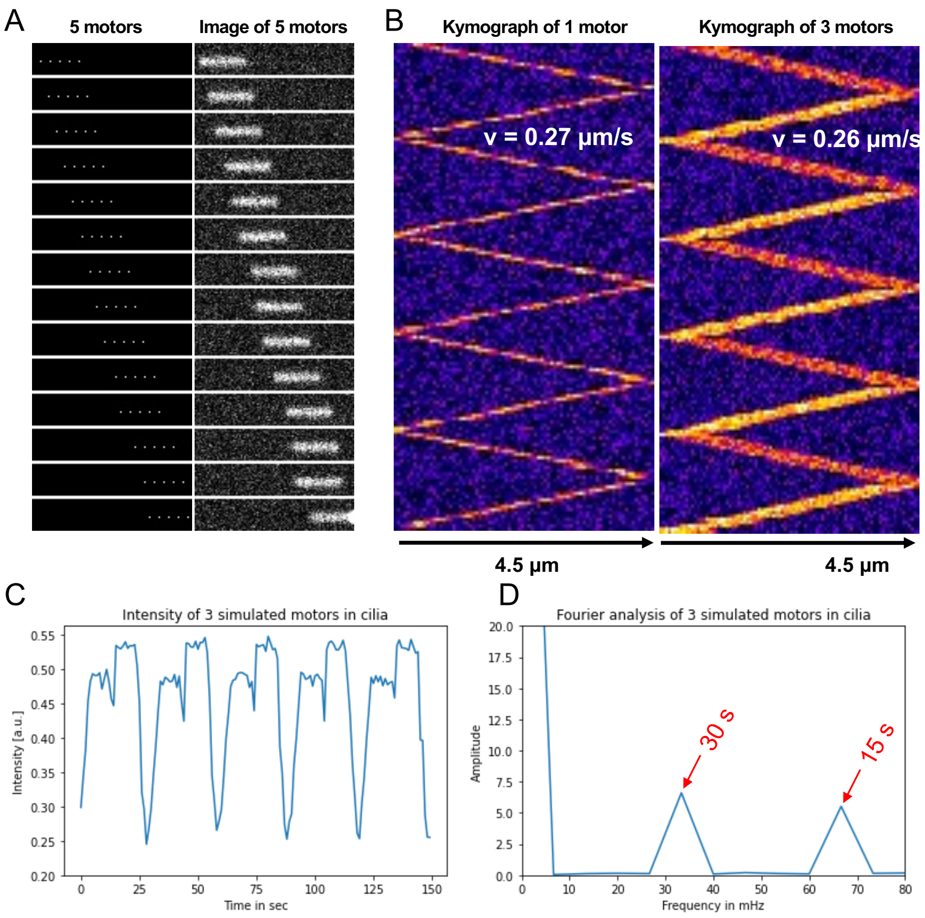


**Figure S2. Image simulations of cilia movement. Related to Figure 1.** (**A**) Left panel shows five simulated motors as consecutive single pixels and right panel shows the appearance of those motors through the microscope. (**B**) Kymographs calculated for images of one or 3 motors were used for validation of the analysis procedure for a nominal velocity of 0.25 µm/sec. The true velocity can be recovered with >95% accuracy. (**C**) Integrated intensity was quantified for a portion of the simulated cilia to emulate time-resolved measurements of eGFP-KIF13B in cilia (compare with Figure 1B, C). There were two periodic intensity peaks, since the quantification of cilia intensity allowed for partial exit of the simulated motors at the cilia base. (**D**) Fourier transform of the intensity profile reveals this 2-fold periodicity, the higher frequency corresponds to one-way transport and the lower frequency (given the periodicity of 30 sec) corresponds to the full bidirectional movement of the particles.

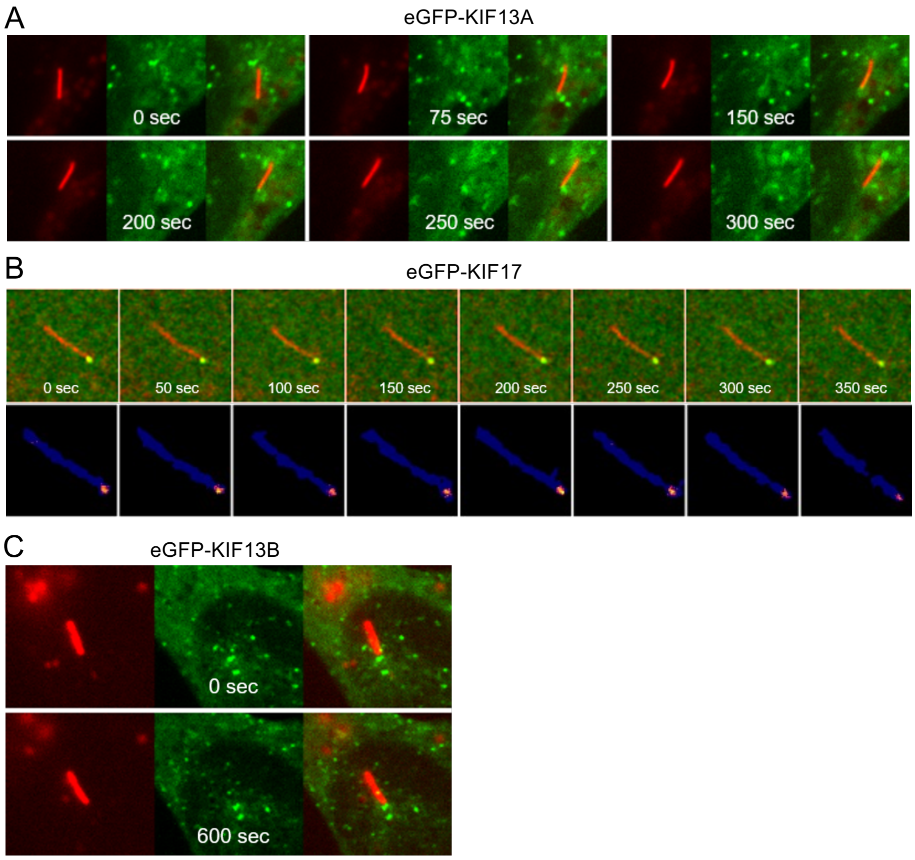


**Figure S3. Control experiments for eGFP-KIF13B live cell imaging analyses.** (**A**, **B**) Frames of representative movies from live cell imaging analysis of eGFP-KIF13A (**A**) and GFP-KIF17 (**B**) in hTERT-RPE1 SMO-tRFP cells. For a quantitative overview of the results, see Table 1. (**C**) Frames of movies from live cell imaging of eGFP-KIF13B in IMCD3 cells stably expressing mCherry-Arl13b (red). In these cells eGFP-KIF13B is localized at the two centrioles at the base of the cilia, marked by mCherry-Arl13b. A total of 10 cells were imaged in the absence of purmorphamine stimulation, and another 20 cells were imaged in the presence of purmophamine, which yielded identical results.

### Supplementary movie legends

**Movie 1. Live cell imaging movie showing transient, burst-like intraciliary movement of eGFP-KIF13B in hTERT-RPE1 SMO-tRFP cells. Related to Figure 1A.** The time lapse video was acquired with a speed of 0.2 frames/second.

**Movie 2. Live cell imaging movie showing transient, burst-like intraciliary movement eGFP-KIF13B in hTERT-RPE1 SMO-tRFP cells. Related to Figure 1D.** The time lapse video was acquired with a speed of 0.3 frames/second.

**Movie 3. Live cell imaging movie showing transient, burst-like intraciliary movement of eGFP-KIF13B in hTERT-RPE1 ARL13B-mScarlet cells. Related to Figure S1D.** The time lapse video was acquired with a speed of 0.9 frames/second.

**Movie 4. Live cell imaging movie of IFT172-eGFP in hTERT-RPE1 cells. Related to Figure 2C, D.** The time lapse video was acquired with a speed of 1.9 frames/second.

**Movie 5. Live cell imaging movie of eGFP-KIF13B cilia localization in Ciliobrevin D-treated hTERT-RPE1 SMO-tRFP cells. Related to Figure 3A, B.** The time lapse video was acquired with a speed of 0.2 frames/second.

**Movie 6. Live cell imaging movie of motorless eGFP-KIF13B in hTERT-RPE1 SMO-tRFP cells. Related to Figure 3C, D.** The time lapse video was acquired with a speed of 0.2 frames/second.
